# MetamORF: A repository of unique short Open Reading Frames identified by both experimental and computational approaches for gene-level and meta-analysis

**DOI:** 10.1101/2020.11.12.380055

**Authors:** Sebastien A. Choteau, Audrey Wagner, Philippe Pierre, Lionel Spinelli, Christine Brun

**Affiliations:** Aix-Marseille Univ., INSERM, TAGC, Turing Centre for Living Systems, Marseille, France; Aix-Marseille Univ., CNRS, INSERM, CIML, Turing Centre for Living Systems, Marseille, France; Institute for Research in Biomedicine (iBiMED) and Ilidio Pinho Foundation, Department of Medical Sciences, University of Aveiro, 3810-193 Aveiro, Portugal; Shanghai Institute of Immunology, School of Medicine, Shanghai Jiao Tong University, Shanghai, China; CNRS, Marseille, France

## Abstract

The development of high-throughput technologies revealed the existence of non-canonical short open reading frames (sORFs) on most eukaryotic RNAs. They are ubiquitous genetic elements highly conserved across species and suspected to be involved in numerous cellular processes. MetamORF (http://metamorf.hb.univ-amu.fr/) aims to provide a repository of unique sORFs identified in the human and mouse genomes with both experimental and computational approaches. By gathering publicly available sORF data, normalizing it and summarizing redundant information, we were able to identify a total of 1,162,675 unique sORFs. Despite the usual characterization of ORFs as short, upstream or downstream, there is currently no clear consensus regarding the definition of these categories. Thus, the data has been reprocessed using a normalized nomenclature. MetamORF enables new analyses at loci, gene, transcript and ORF levels, that should offer the possibility to address new questions regarding sORF functions in the future. The repository is available through an user-friendly web interface, allowing easy browsing, visualization, filtering over multiple criteria and export possibilities. sORFs could be searched starting from a gene, a transcript, an ORF ID, or looking in a genome area. The database content has also been made available through track hubs at UCSC Genome Browser.

## INTRODUCTION

Short open reading frames (sORFs) are usually defined as sequences delimited by a start and a stop codon and potentially translatable into proteins of less than 100 amino acids (1–8). They are present on all classes of transcripts (including presumptive long non-coding RNAs) and have been identified on most eukaryotic RNAs (2, 5, 8–15). In addition, their sequence often begin with a non-canonical start codon (8). Consequently, they have long been overlooked and interest in their possible regulatory functions has only raised recently with the advent of the ribosome profiling method that strongly suggests their translation (1, 3, 5, 6, 16–22).

Several sORFs biotypes have been defined according to their location on RNAs. For instance, upstream ORFs (uORFs) are located in the 5’ untranslated regions of mRNAs and have been defined as sORFs whose start codon precedes the main coding sequence (CDS) (6, 8, 17, 18, 23). They are conserved across species (5, 6, 11, 21, 24), although they seem to be less conserved than canonical protein-coding ORFs (25). To date, uORFs have been essentially reported as gene expression *cis*-regulatory elements that regulate the efficiency of translation initiation of the main CDS, notably alleviating the repression of translation during cellular stress (13, 17, 18, 20, 23, 26). Moreover, the discovery of uORF-, and more generally sORF-encoded peptides led to the assumption that they may also play functional roles in *trans* (2–4, 7, 9, 10, 18, 24, 27–30), for instance as ligands of major histocompatibility complex (MHC) molecules (12, 22, 23). Very interestingly, uORF-encoded peptides have also been shown to form protein complexes with the protein encoded by the main CDS of the same mRNA (31) and it has been suggested that polycistronic sequences may exist in Eukaryotes (24, 31). Furthermore, given the increasing evidence on the regulatory functions of peptides encoded by sORFs located within mRNAs, introns of pre-mRNAs, lncRNAs, primary transcripts of miRNAs or rRNAs (2, 8–15, 26), there is an urgent need to study sORFs *(i)* individually, and *(ii)* at the whole proteome scale. Indeed, the latter should reveal important sORFs features, thus enabling the characterization and the identification of their functions. However, the fact that *(i)* the publicly available data are scattered across different databases and *(ii)* datasets are aligned on different genome builds, differently annotated and formatted, calls for an uniformized resource where each sORF is individually described. With this in mind, we have built a resource database of publicly available sORFs identified in the human and mouse genomes, by gathering information from computational predictions, Ribo-seq and proteomic experiments. The curation of data, their homogenization in order to merge the redundant information into unique entries, the completion and computation of missing information (*e.g*., sequences, Kozak contexts) and the reannotation of sORF classes represent the added value of this database. Notably, this enables the analysis at locus, gene, transcript, and ORF levels, as well as groups of them. In this work, we propose *(i)* a pipeline to regularly update the content of the database in a reproducible manner, *(ii)* a database content that can be fully downloaded for custom computational analyses and *(iii)* a user-friendly web interface to ease data access to biologists.

## MATERIAL AND METHODS

### MetamORF pipeline and database development

#### Inclusion criteria for publicly available sORF-related data

A total of 18 data sources, either *H. sapiens* and *M. musculus* original datasets or re-processed publicly available sORFs repositories, have been considered for inclusion in our database (Supplementary Table S1) (5, 7, 11, 12, 14, 15, 17–22, 32–37). These data sources provide results from computational predictions, Ribo-seq experiment analyses and mass spectrometry (proteomics / proteogenomics) analyses. The data sources not providing the absolute genomic coordinates of the ORF start and stop codons (5, 17, 20, 32–34) or fully included in another data source considered here (21), have been discarded. Databases that did not allow export of their content in a single file or to automate the download of all the files from their website, have also been discarded (19, 35). Despite their short size, it has been noticed that sORFs can be spliced. Theoretical lengths of the ORFs have been computed as the distance between the start and stop codons, eventually removing the intron length(s) when information about the ORF splicing was provided. Due to splicing, the theoretical length and the one reported by the data source may be different. Data sources harboring this difference for more than 95% of their entries were discarded as this indicates the splicing information was missing (10). Finally, data sources for which we were not able to perform this assessment as they were not providing information regarding the splicing of the ORF and did not provide any ORF length (15, 36) have not been included as well. Hence, the database has been made by collecting data from six distinct sources (Figure 1), including either original datasets (Table 1 and Supplementary Table S2) (11, 12, 14, 18, 22) or reprocessed data (37), and discarding 12 of them (Supplementary Table S1). Notably, we have included data from sORFs.org (37), considered as the main and most comprehensive repository of sORFs identified by genome-wide translation profiling (Ribo-seq), that currently integrates re-processed data from 73 original publications.

**Figure 1.**
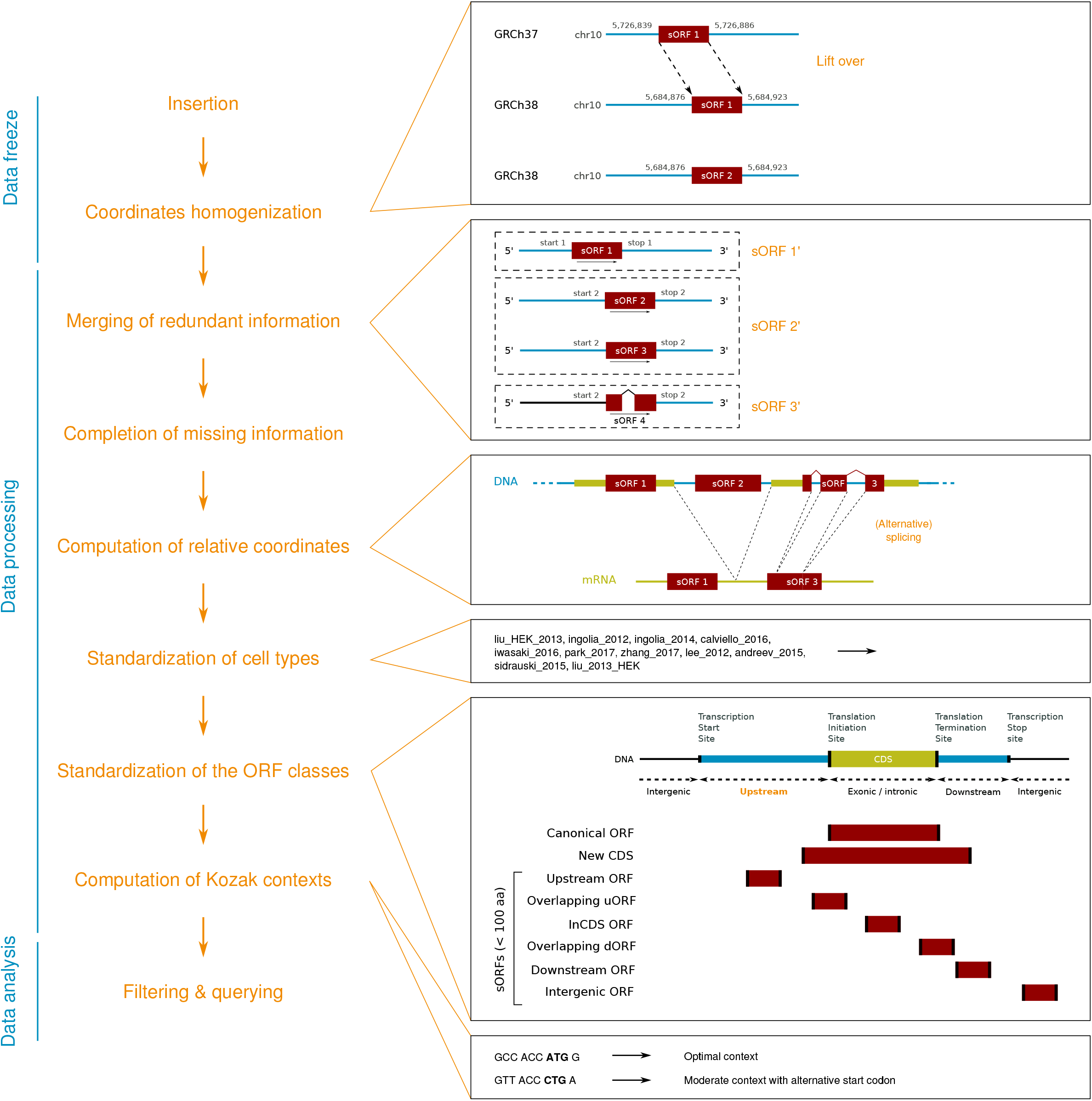
MetamORF pipeline. This figure represents the workflow used to build MetamORF. First, the data from the sources selected has been inserted in the database and the absolute genomic coordinates have been homogenized from their original annotation version to the most recent (GRCh38 or GRCm38). Then the redundant information, *i.e*. the entries describing the same ORFs (same start, stop and splicing), have been merged, allowing to get one single and unique entry for each ORF detected on the human and mouse genomes. The missing information (sequences, transcript biotype) have been downloaded from Ensembl and the ORF relative coordinates have been computed. Finally the cell types and ORF classes have been normalized and the Kozak contexts have been computed using the sequences flanking the start codons.

**Table 1.** Information about the data sources used to build MetamORF. See supp. table S1 for more information about these data sources.

| Publication | DOI |
| --- | --- |
| Mackowiak et al., 2015, Genome Biol. (11) | 10.1186/s13059-015-0742-x |
| Erhard et al., 2018, Nat. Meth. (22) | 10.1038/nmeth.4631 |
| Johnstone et al., 2016, EMBO (18) | 10.15252/embj.201592759 |
| Laumont et al., 2016, Nat. Commun. (12) | 10.1038/ncomms10238 |
| Samandi et al., 2017, eLife (14) | 10.7554/eLife.27860 |
| Olexiouk et al., 2018, Nucl. Ac. Res. (37) | 10.1093/nar/gkx1130 |

For each of these sources, a set of features essential to properly characterize the sORFs, related to their location, length, sequences, environmental signatures and cell types (*i.e*. cell lines, tissues or organs) in which they are expressed, have been collected (see Table 2 for a full list of features considered for inclusion). When it was not provided by the source, the symbol of the gene related to the sORF was recovered using the transcript identifier (ID, if provided) or searching for the gene(s) or ncRNA(s) overlapping with the sORF coordinates in the original annotation version by querying Ensembl databases in their appropriate versions (v74, 75, 76, 80, 90) with pyensembl (v1.8.5, https://github.com/openvax/pyensembl). In addition to these features, information regarding the transcript(s) harboring the ORFs have been collected from the data sources when available. This is of particular interest as some ORF features, such as the ORF class, may depend on the transcript they are located on (*e.g*. an ORF may be located in the 5’UTR of a transcript and be overlapping with the CDS of another transcript). Finally, 3,379,219 and 2,066,627 entries from these six data sources have been collected and inserted in MetamORF for *H. sapiens* and *M. musculus*, respectively (Table 3).

**Table 2.**
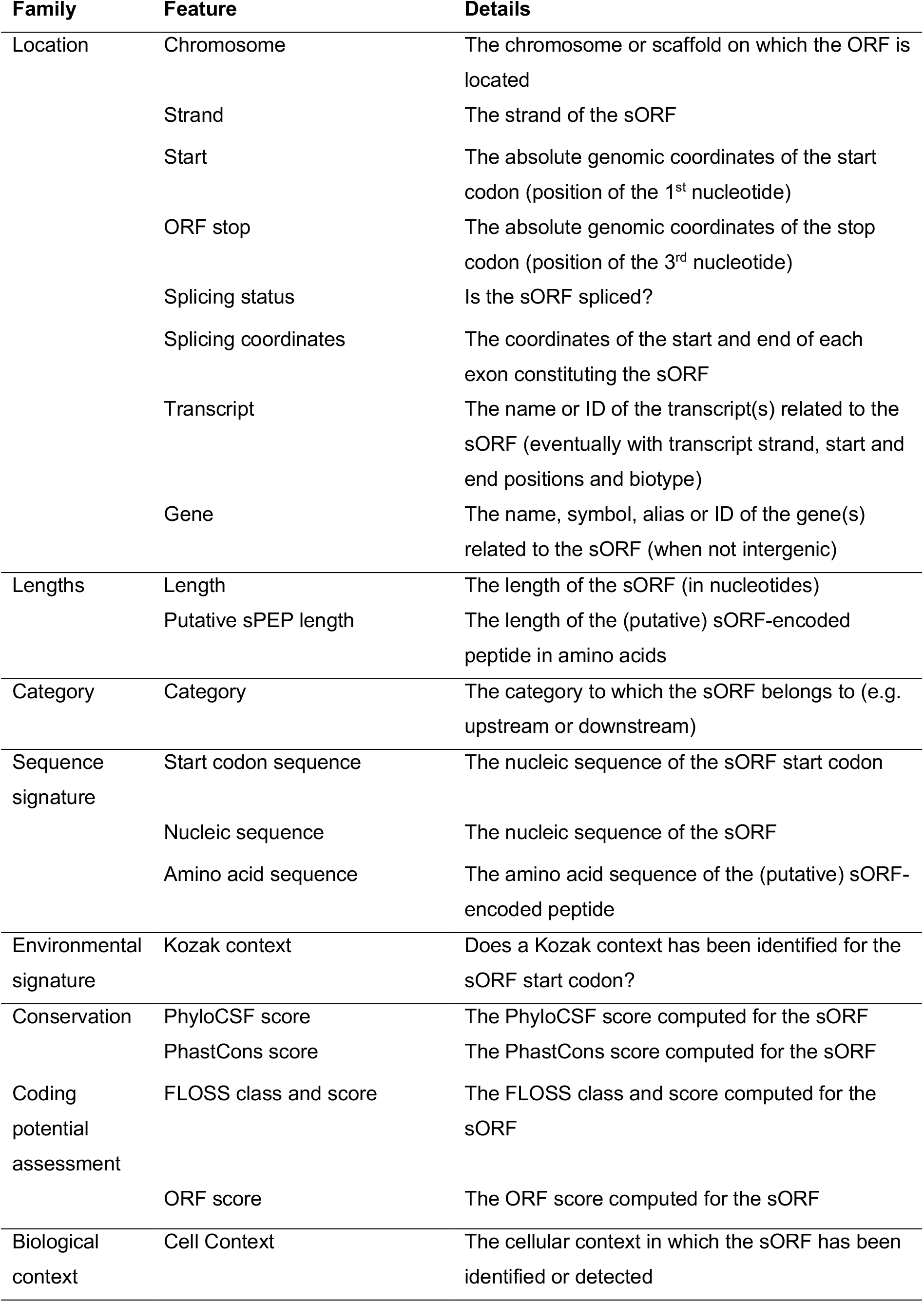
Features allowing to characterize the sORFs.

| **Family** | **Feature** | **Details** |
| --- | --- | --- |
| Location | Chromosome | The chromosome or scaffold on which the ORF is located |
|  | Strand | The strand of the sORF |
|  | Start | The absolute genomic coordinates of the start codon (position of the 1<sup>st</sup> nucleotide) |
|  | ORF stop | The absolute genomic coordinates of the stop codon (position of the 3<sup>rd</sup> nucleotide) |
|  | Splicing status | Is the sORF spliced? |
|  | Splicing coordinates | The coordinates of the start and end of each exon constituting the sORF |
|  | Transcript | The name or ID of the transcript(s) related to the sORF (eventually with transcript strand, start and end positions and biotype) |
|  | Gene | The name, symbol, alias or ID of the gene(s) related to the sORF (when not intergenic) |
| Lengths | Length | The length of the sORF (in nucleotides) |
|  | Putative sPEP length | The length of the (putative) sORF-encoded peptide in amino acids |
| Category | Category | The category to which the sORF belongs to (e.g. upstream or downstream) |
| Sequence signature | Start codon sequence | The nucleic sequence of the sORF start codon |
|  | Nucleic sequence | The nucleic sequence of the sORF |
|  | Amino acid sequence | The amino acid sequence of the (putative) sORF-encoded peptide |
| Environmental signature | Kozak context | Does a Kozak context has been identified for the sORF start codon? |
| Conservation | PhyloCSF score | The PhyloCSF score computed for the sORF |
|  | PhastCons score | The PhastCons score computed for the sORF |
| Coding potential assessment | FLOSS class and score | The FLOSS class and score computed for the sORF |
|  | ORF score | The ORF score computed for the sORF |
| Biological context | Cell Context | The cellular context in which the sORF has been identified or detected |

**Table 3.**
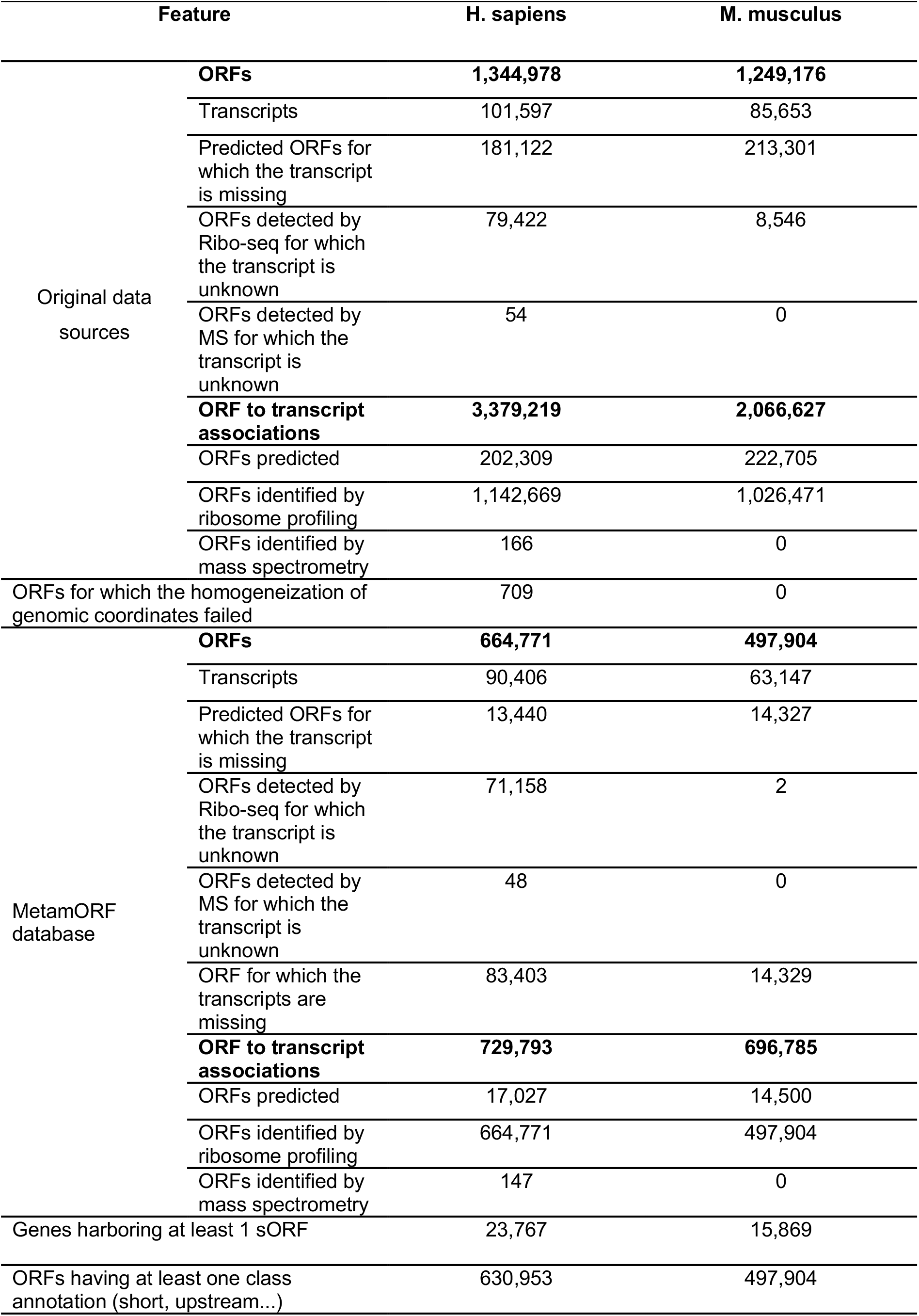
MetamORF most important statistics.

#### Homogenization of genomic coordinates

As the data sources were providing genomic coordinates from different genome annotation versions (*e.g*. GRCh38 and GRCh37), all the genomic coordinates registered in our database have been lifted over the latest annotation version (GRCh38 for *H. sapiens*, GRCm38 for *M. musculus*) using pyliftover (v0.4, https://pypi.org/project/pyliftover). The lift-over has been considered as failed for an entry if *(i)* at least one of the coordinates (*i.e*. start, stop or one of the start or end exon coordinates) was located on a strand different from all the others, or *(ii)* the chromosome of the position changed during the lift-over, or *(iii)* the distance (in nucleotides) between the sORF start and stop codons has changed after the lift-over. All the entries for which the lift-over failed were removed from the database. Based on the previous assumptions, the lift-over failed for 709 ORFs (377 failed due to the last criteria) in *H. sapiens* and for none of the *M. musculus* entries (Table 3). The choice of such stringent criteria has been strengthened by the fact that these entries only represent less than 0.05% of the total number of entries for *H. sapiens* and are more susceptible to be unreliable entries.

#### Merge of redundant information

As our database aims to provide a repository of unique identified sORFs of the human and mouse genomes, all the redundant entries describing the same sORFs have been merged. In a first step, we identified all the sORF entries for which all the identification features were provided (chromosome, strand, start position, stop position, splicing status and splicing coordinates). sORFs sharing the same feature values were merged. In a second step, we identified all the remaining entries with only partial identification features provided: the chromosome as well as either *(i)* both the strand and the start position, or *(ii)* both the strand and the stop position, or *(iii)* both the start and the stop position. Those entries were merged to the best matching fully described entries identified in the first step. If no matching fully described entry was found, then the entries were removed. In order to keep track of the number of times a same sORF has been described in the original data sources, the initial number of entries merged together was registered for each sORF.

During this merging, information regarding the transcripts that harbor the sORFs have been registered too. Hence, when several sORFs were merged into one single entry in MetamORF, the resulting new sORF entry was registered as harbored by all the distinct transcripts related with the original entries. After this removal of redundant information, we were finally able to identify 664,771 and 497,904 unique sORFs for *H. sapiens* and *M. musculus*, respectively (Table 3).

It should be noticed that all unique sORF entries generated at this stage have been kept, including the ones describing ORFs longer than 100 amino acids. Entries describing such ORFs may be either coming from data sources that *(i)* did not remove the ORFs longer than 100 amino acids, or *(ii)* used a higher threshold or *(iii)* described the ORF as unspliced whilst it is actually susceptible to be spliced (and thus has a shorter sequence on the transcript than the one expected).

#### Completion of missing information and computation of relative coordinates

In the original data sources, the only information provided (when provided), on the transcripts was the transcript ID. Detailed information was retrieved from Ensembl databases (v90) through their REST API and inserted in our database: *(i)* the biotype, *(ii)* the transcript start and end genomic coordinates, *(iii)* the codon of the canonical coding sequence (CDS, for protein-coding transcripts only) start and stop genomic coordinates and *(iv)* the full nucleic sequence. In addition, the sequence flanking the start codon (20) has been recovered. As the sORF nucleic and amino acid sequences were not systematically provided by the data sources, these were downloaded from the Ensembl databases using their genomic coordinates.

Moreover, when the transcript ID was available, sORF start and stop relative coordinates have been computed on each of their transcript using AnnotationHub (v2.18.0, (39)) and ensembldb (v2.10.2, https://bioconductor.org/packages/release/bioc/html/ensembldb.html) R packages (R v3.6.0).

#### Standardization of the cell types and ORF classes

##### Cell types

Original data sources do not use a common thesaurus or ontology to name the cell types (*e.g*., ‘HFF’ and ‘Human Foreskin Fibroblast’) or use non-biological meaning names (*e.g*., sORFs.org (37) provides the name of the original publication as a cell type). In order to provide an uniform informative naming, we manually recovered the name of the cell line, tissue or organ used in these datasets and defined an unique name to be used in our database for each cell line, tissue or organ (Supplementary Table S3).

##### ORF classification

Despite the use of a common nomenclature by the wide majority of the scientific community to annotate the open reading frames, based on their relative position on their transcript (*e.g*., short, upstream, downstream, overlapping), no clear consensus about the definitions of these categories nor their names has been defined so far (25). In order to homogenize this information in MetamORF, we created a new annotation of the ORFs using both the ORF length, transcript biotype, relative positions and reading frame information when available. In this annotation, a threshold of 100 amino acids has been used to define the “short ORFs”, as this value is the most commonly used for historical reasons (2, 4, 6, 8, 24).

#### Computation of the Kozak contexts

The Kozak motif and context have been regarded as the optimal sequence context to initiate translation in all eukaryotes. We have thus assessed the Kozak context for each sORF, using the criteria defined in Hernandez et al. (40). Briefly, for each ORF to transcript association, the Kozak context was computed looking for regular expression characterizing an optimal, strong, moderate or weak Kozak context (Supplementary Tables S4 and S5). Kozak-alike contexts were also computed for non-ATG initiated sORFs looking for the same patterns with flexibility regarding nucleotides at +1 to +4 positions.

### MetamORF softwares and languages

The pipeline used to build MetamORF has been developed using Python (v2.7) with SQLAlchemy ORM (sqlalchemy.org, v1.3.5). The database has been handled using MySQL (mysql.com, v8.0.16). Docker (docker.com, v18.09.3) and Singularity (singularity.lbl.gov, v2.5.1) environments have been used in order to ensure reproducibility and to facilitate deployment on high-performance clusters (HPCs).

The MetamORF web interface has been developed using the Laravel (laravel.com, v7.14.1) framework with PHP (v7.3.0), JavaScript 9, HTML 5 and CSS 3. The NGINX (v1.17.10) web server PHP server (v7.3.0) were deployed with Docker (docker.com, v18.09.3) and Docker-compose (v1.24.0) to ensure stability.

## DATABASE CONTENT, ACCESSIBILITY AND WEB INTERFACE

### A new repository of short ORF-related data

MetamORF describes 664,771 and 497,904 unique ORFs in the human and mouse genomes respectively, providing at least the information necessary to locate the ORF on the genome, its sequence and the gene it is located on (excepted for intergenic ORFs). Extensive information related to the transcripts is provided respectively for 614,997 (~93%) and 497,904 (100%) sORFs for the human and mouse genomes respectively. These features allowed us to classify 630,953 (~95%) human ORFs and 497,904 (100%) mouse ORFs in at least one class (Table 3, Figure 2, Supplementary Figure S1). Interestingly, it should be noticed that a large proportion (36% and 52% respectively for *H. sapiens* and *M. musculus*) of ORFs are using an alternative frame to the main CDS. In addition, nearly 23% of the ORFs are located on non-coding RNAs for both species.

**Figure 2.**
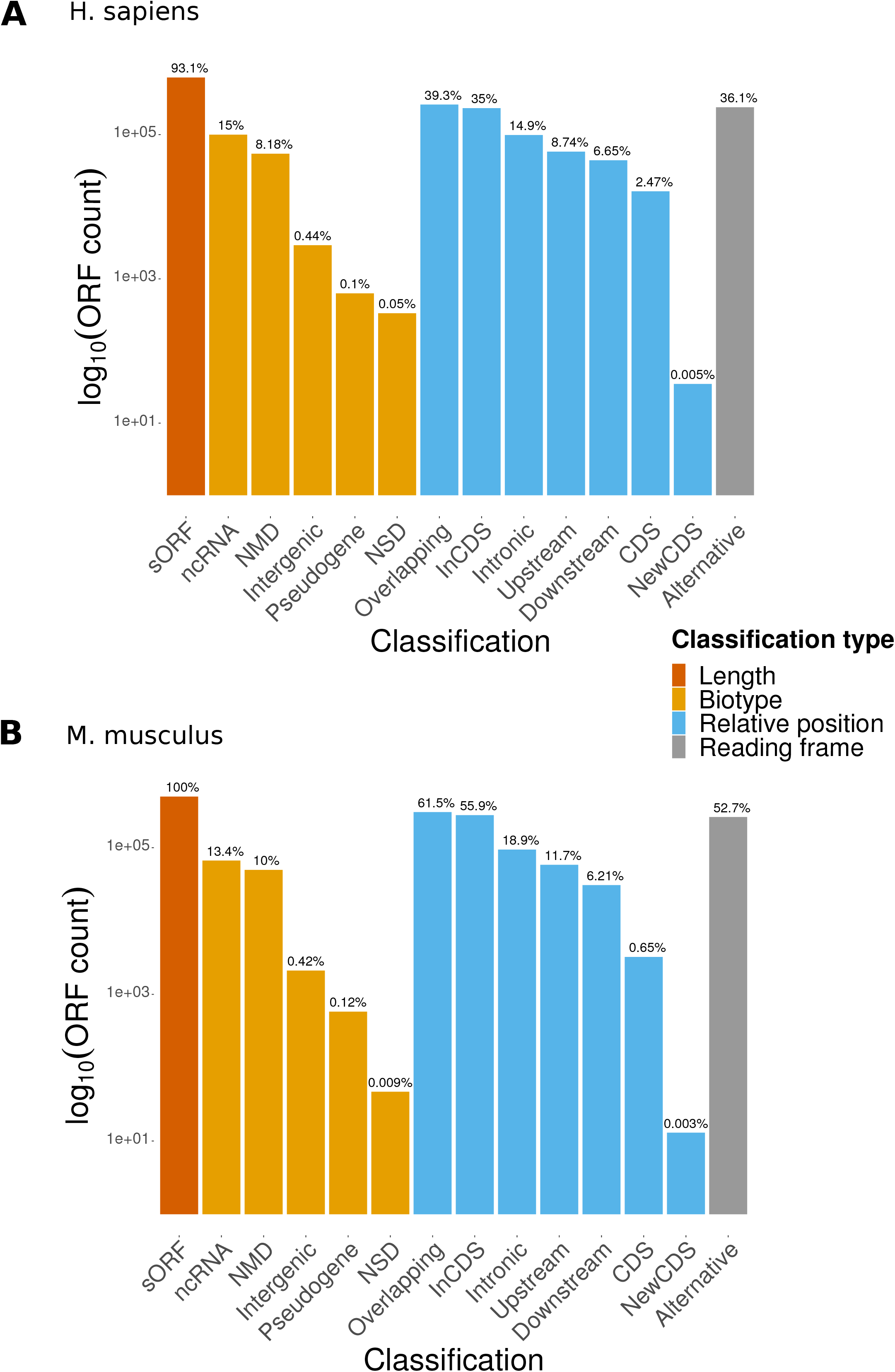
Count of ORFs in each class. The barplot represent the count of ORFs annotated for each class for (A) *H. sapiens* and (B) *M*. musculus. The percentages displayed over the bars indicates the proportion of ORFs annotated in the class over the total number of ORFs registered in the database for the species.

### User-friendly web interface and genome tracks

To provide users with a clear, fast and easy-to-use database, MetamORF can be queried through an user-friendly web interface at http://metamorf.hb.univ-amu.fr. A tutorial as well as a documentation page are available online. Briefly, the users may search for sORFs contained in the database starting with a gene symbol (symbol, alias, ID), transcript ID (ID, name), ORF ID, or screening a particular genomic area. The data is made accessible through four types of pages: *(i)* a “gene-centric” page (Figure 3), allowing to visualize information related to all transcripts and sORFs on a gene, *(ii)* a “transcript-centric” page, allowing to browse information related to a transcript gene and all its sORFs, *(iii)* an “ORF-centric” page allowing to fetch information related to all transcripts and gene that harbor the chosen ORF and finally *(iv)* a “locus” page allowing to get information related to all sORFs located in a particular locus. It is possible to navigate from one to another page easily to get extensive information about an sORF, a gene or a transcript (Supplementary Figure S2).

**Figure 3.**
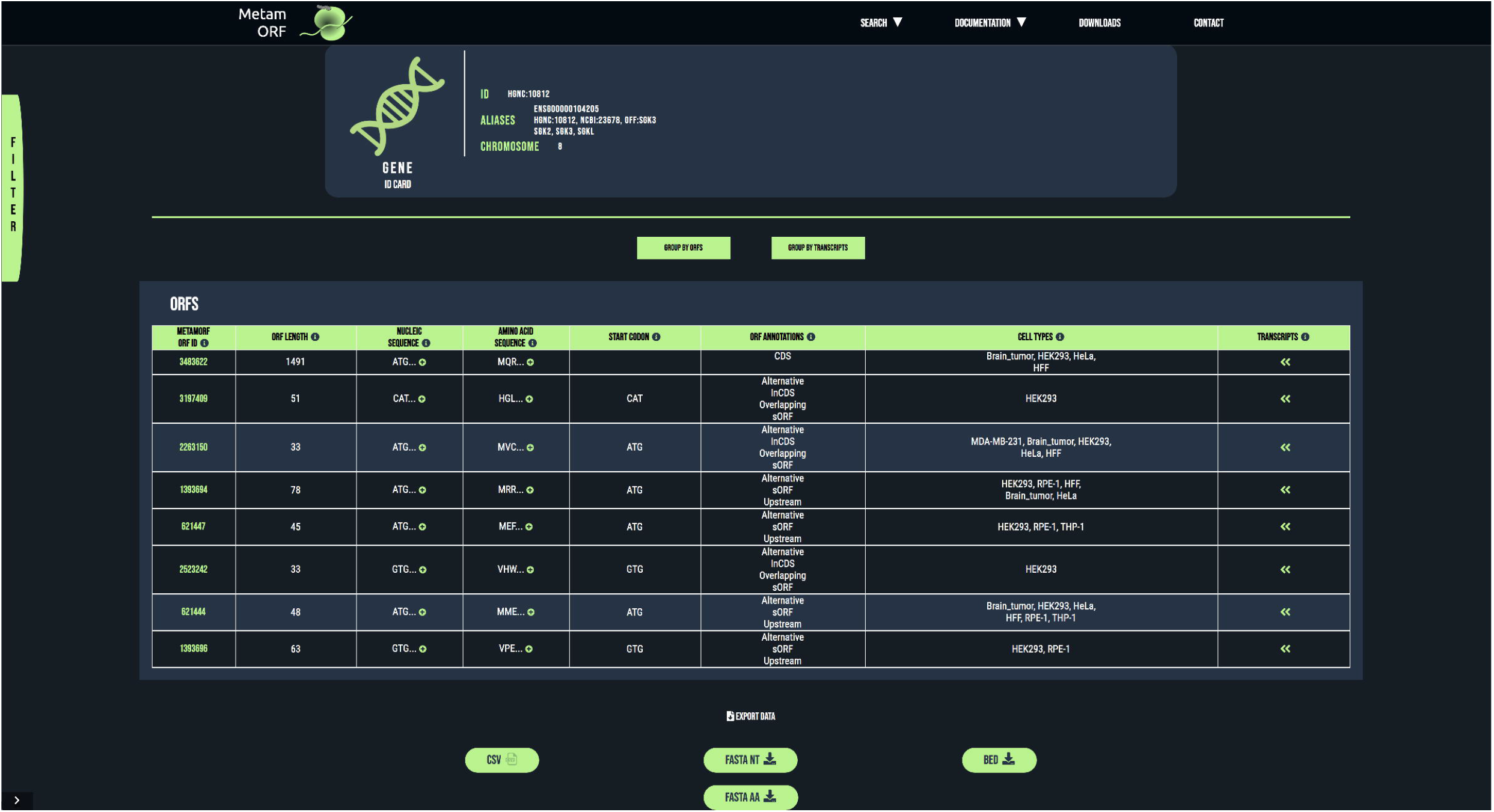
MetamORF gene-centric view. The page displays the transcripts and the ORFs related to SGK3 gene. A filter has been applied to select exclusively the ORFs detected in HFF, Jurkat, RPE-1, HEK293 or HeLa cells. Other filters may be used and the results can be exported as CSV, FASTA or BED files.

In each page, the results can be filtered on: *(i)* the identification method (computational prediction, ribosome profiling or mass spectrometry), *(ii)* the start codon, *(iii)* the Kozak context (as previously defined), *(iv)* the genomic length (defined as the sum of lengths of each exon constituting the ORF), *(v)* the transcript biotype (according to the Ensembl definitions), *(vi)* the ORF annotation (as previously defined) and *(vii)* the cell type (Supplementary Table S3).

All results can be exported in an easily-parsable format (comma-separated values file, CSV), as well as in FASTA or BED format.

On ORF, transcript and locus pages, a link allowing to easily visualize all the ORFs localized in a particular area on the UCSC genome browser (41) is proposed. We also implemented genome track hubs, allowing to use UCSC genome browser advance options, such as filtering on ORF categories, transcript biotypes, cell types and transcript IDs.

In addition to this user-friendly interface, it is possible to download from the website the content of the full MetamORF database at BED, and FASTA formats.

## DISCUSSION AND CONCLUSION

MetamORF contains data about 1,162,675 unique sORFs for the human and mouse genomes identified by both experimental and computational approaches. Whilst the Ribo-seq is considered by most as the “gold standard” method to identify sORFs experimentally, the added value of predictive computational approaches, proteomics and peptidomics to characterize such biological sequences remains certain. Because these technologies are offering complementary information at genomic, transcriptomic and proteomic scales, we decided to include data from both experimental and computational experiments in our database. Nevertheless, data coming from distinct data sources may be difficult to compare, in particular because they are not necessarily using the same genome annotation and definitions of ORF classes and Kozak contexts, for instance. By homogenizing this information, MetamORF offers the possibility to compare datasets coming from different sources. We noticed that information regarding the Kozak context is missing most of the time and start flanking sequences are usually not provided. Hence, MetamORF provides there a new interesting set of information.

It should be noticed that a large amount (~80 %) of the sORFs contained in our database have been described in the sORFs.org repository (37). Despite being the most prominent sORF database and offering the community data processed in a normalized way using their own workflow, it has already been highlighted that sORFs.org does not provide metagene analyses (1). In addition, such analysis is made difficult by the absence of gene names as well as the high redundancy of information contained in the sORF.org database (37), an issue we addressed with MetamORF. Hence, in comparison with existing resources, MetamORF allows analyses at ORF, transcript, gene and loci levels. In addition, it opens the possibility of studying sORFs as a group, at a global scale.

The resource is accessible at http://metamorf.hb.univ-amu.fr and provides an intuitive querying interface to enable wet lab researcher to easily question this large set of information. Moreover, the implementation of MetamORF content in track hubs allows both quick and advanced visualization of data through the UCSC genome browser. Finally, the database content may be exported at various convenient formats widely used by the scientific community (e.g. FASTA, BED).

We believe that MetamORF is of interest not only to bioinformaticians working on short ORFs but also to a wider community, including any biologist who may benefit from knowledge regarding the sORFs located on their gene, transcript or region of interest. As ribosome profiling becomes more appreciated and proteomics starts to allow accurate identification of short peptides, new data describing sORFs in various conditions are expected to be published in the next years, and our database is expected to grow accordingly. As a conclusion, we believe that MetamORF should help to address new questions in the future, in particular regarding the regulatory functions of the sORFs as well as the functions of the short peptides they may encode.

## DATA AVAILABILITY

Data sources are available on the editor’s website or using the links provided in their original publications. The source code used to create the database, and the full technical documentation (source code documentation, manual, database structure, dockerfiles) are available on GitHub (https://github.com/TAGC-NetworkBiology/MetamORF). Full content of the database can be downloaded at BED and FASTA formats from MetamORF website and up-to-date version of track hubs may be download and/or used with your favorite genome browser providing the following link: http://metamorf.hb.univ-amu.fr/hubDirectory/hub.txt. The dump of the database is available on request.

## SUPPLEMENTARY DATA

Supplementary Data are available upon request.

## ACKNOWLEDGEMENT

We thank Andreas Zanzoni for helpful discussions.

